# ERK Allosteric Activation: The Importance of Two Ordered Phosphorylation Events

**DOI:** 10.1101/2025.02.27.640630

**Authors:** Clil Regev, Jang Hyunbum, Ruth Nussinov

## Abstract

ERK, a coveted proliferation drug target, is a pivotal kinase in the Ras/ERK signaling cascade. Despite this, crucial questions about its activation have not been fully explored on the foundational, conformational level. Such questions include (i) *Why ERK’s activation demands dual phosphorylation*; (ii) *What is the role of each phosphorylation site in the activation loop*; and (iii) *Exactly how the (ordered) phosphorylation steps affect the conformational ensembles of the activation loop*, *their propensities and restriction to a narrower range favoring ERK’s catalytic action*. Here we used explicit molecular dynamics simulations to study ERK’s stability and the conformational changes in different stages along the activation process. The initial monophosphorylation event elongates the activation loop to enable the successive phosphorylations, which reintroduce stability/compactness through newly formed salt bridges. The interactions formed by the monophosphorylation are site-dependent, with threonine’s phosphorylation presenting stronger electrostatic interactions compared to tyrosine’s. Dual phosphorylated ERKs revealed a compact kinase structure which allows the HRD catalytic motif to stabilize the ATP. We further observe that the hinge and the homodimerization binding site responded to a tri-state signaling code based solely on the phosphorylation degree (unphosphorylated, monophosphorylated, dual phosphorylated) of the activation loop, confirming that the activation loop can allosterically influence distant regions. Last, our findings indicate that threonine phosphorylation as the second step is necessary for ERK to become effectively activated and that activation depends on the phosphorylation order. Collectively, we offer ERK’s dual allosteric phosphorylation code in activation and explain why the phosphorylation site order is crucial.

## Introduction

The mitogen-activated protein kinase (MAPK) pathway is among the most highly conserved signaling cascades in the cell [1], controlling cell proliferation, differentiation, and survival as a response to external stimuli of growth factors [1–3]. MAPK’s cascading activation events involve four proteins: a GTPase, Ras, and three kinases; Raf, mitogen-activated protein kinase kinase (MEK), and its end node, extracellular signal-regulated kinase (ERK). Among the four, ERKs are the only components not confined to the cytoplasm, but also translocating to the nucleus. ERKs activate diverse substrates, including cytoskeletal proteins and transcription factors, governing gene expression, cytoskeletal reorganization, and cellular growth [4–7]. Because of their broad functionality, ERKs activity is highly regulated through subcellular localization [6], coupling with phosphatases [7, 8], scaffolding proteins [9, 10], negative feedback loops [11], and diffusion that relies on dimerization [12–14]. Their dysregulation can result in diverse cancers, including melanoma, colorectal, and lung cancer [15].

The sequences of ERK’s common isoforms, ERK1 (MAPK3) and ERK2 (MAPK1) are 84% identical, with longer N- and C-terminal tails for ERK1 [16, 17]. Like many kinases, ERK2 (hereafter ERK) structure is bilobed, with an N-terminal lobe connected to a larger C-terminal lobe by a flexible hinge region (**Figure S1A**). The N-lobe contains a single α-helix (αC-helix) and a β-sheet. The β-sheet’s glycine-rich sequence forms a loop (G-loop) that binds the αC-helix via a Lys-Glu salt bridge (**Figure S1B**). The C-lobe consists primarily of α-helices with a few flexible segments. These segments include the activation loop at the front top of the C-lobe, a hinge in the back interface of the lobes, and a section preceding the last L16 segment containing the α_L_16-helix. Unlike other kinases, the C-lobe’s terminal α_L_16-helix interacts with the N-lobe and binds the αC-helix, limiting lobes dissociation, and restricting ERK’s dimensions and conformations.

The ATP binding site is in the cleft between the two lobes, underneath the G-loop. Regulatory sites such as the DFG domain can stabilize the active conformation by forming supportive spines along the lobes, and the HRD catalytic segment stabilizes the substrate complex prior to deprotonation [18]. Based on experimental structures and simulated models, ERK’s active and inactive states are similar for these domains, making the conformation of the activation loop and the residues orientation within the ATP pocket key factors in distinguishing between the active and inactive states [19].

ERK’s activation loop requires dual phosphorylation by upstream MEKs to become active. These phosphorylations take place on tyrosine and threonine residues in the activation loop, often referred to as the TxY motif. Substituting the phosphate groups on these residues induces changes in the kinase conformation and surface charge that ultimately switch the protein between the inactive and active states [20]. Multiple phosphorylation sites can complexify the kinase activation mechanism, for example, having one site influencing the position of another [21], thus providing the kinase with an additional layer of self-regulation.

Previous studies provided insights into the dual phosphorylation of ERK and its monophosphorylated intermediates [22–24], challenging the common assumption that MEK phosphorylates ERK in a random order. While MEK exhibits equal phosphorylation rates for the sequential phosphorylation of both tyrosine and threonine, the key finding was the identification of tyrosine as the primary phosphorylation site, revealing ERK’s intrinsic selectivity. Studies even suggested that monophosphorylated threonine can be formed only as a hydrolyzed product of dual phosphorylated ERK. These observations were supported by mathematical models [22], and carefully controlled kinetic measurements [23, 24]. The kinetic studies also showed that the second phosphorylation step is twice as slow as the initial monophosphorylation, regardless of whether the tyrosine or threonine was already phosphorylated [23], revealing another layer of complexity in this critical cellular process.

Ongoing efforts aim at elucidating the reasons for kinase hyperactivity. Structural studies focus on allosteric mechanisms, expressed by conformational and dynamic changes, governing regulatory processes like activation and autoinhibition [25–34]. It is common to think of a two-conformation model to explain how phosphorylation activates the protein. In such a scenario, the introduction of phosphate groups locks the loop in a conformation that facilitates the binding of ATP and substrate. However, this simplified model fails to fully describe the transition of ERK from the inactive to the active state due to the conformational similarity between the two states. Instead of the classical two separated conformations, the accepted model for ERK is based on the conformational flexibility especially of the activation loop, with ensembles of conformations [35]. The transition between them is inevitable but phosphorylation can allosterically regulate their equilibrium through strengthened interactions, and drive ERK to populate the preferred state, a concept that was previously demonstrated for effectors [35–39].

The two-ensembles model provides a new perspective and highlights important domains and regions in the kinase that can control the equilibrium. For example, oncogenic mutations alter the conformations of residues within the ATP-binding pocket and the activation loop, resulting in resistance to inhibitors. They work by shifting the ensemble, toward the stabilized active state [33, 40, 41]. Also, competitive inhibitors that bind dual phosphorylated ERK while presenting binding-patterns of an unphosphorylated ERK are more efficient [19, 42]. *In silico* studies reveal the relationship between distant regions of the kinase and the active site, contributing to its fine-tuned regulation [43]. Overall, conformational changes are largely confined to the flexible domains of ERK: the flexible hinge that locks the ATP between the two lobes and the activation loop that cooperatively adapts to different conformations [9, 10].

Here, we focus on the dual phosphorylation and the activation mechanism of ERK at the atomic level by investigating the structural differences of the intermediates and their stability. Our molecular dynamics (MD) simulations sample the conformational ensembles of the inactive and active states. The conformations, residing in basins separated by relatively small free energy differences and low kinetic barriers, have propensities that determine the enzyme’s functional state [39]. We will recount the steps of processive dual phosphorylation on the conformational level, a major component of ERK activation. We will also offer answers to crucial questions such as (i) *Why ERK’s activation demands dual phosphorylation*, (ii) *What is the role of each phosphorylation site in the activation loop*, and (iii) *Does the order of phosphorylation matter in terms of activation*. It is important to note that our computational findings agree with earlier reported experimental observations and provide a rational structural mechanism to explain the need for dual phosphorylation activation.

Taken together, these findings provide a more detailed understanding of the ERK phosphorylation mechanism and may contribute to allosteric drug targeting during the process ERK regulation.

## Results

### Structural compactness is governed by dual phosphorylation

Active kinase conformation is typically described as a closed and compact structure [44]. Studies revealed that by adopting such a conformation, ERK can enhance its interactions with the substrate and ATP [45]. The active, closed conformation is stabilized by two molecular spines (catalytic spine, C-Spine; and regulatory spine, R-spine) that hold the lobes together and minimize their dissociation by the interactions between the N-terminal and C-terminal lobes. To elucidate the structural basis of ERK during activation, we performed MD simulations on ERK with different phosphorylation codes: monophosphorylation of ERK at T185 and Y187 in the activation loop (denoted as ERK^pT^ and ERK^pY^, respectively) and dual phosphorylation of ERK at both residues (denoted as ERK^pTpY^, ERK^pYpT^, and ERK^p(TY)^) (**Figure 1 and Table S1**). ERK^pTpY^ and ERK^pYpT^ were subjected to sequential phosphorylation at each monophosphorylated ERK during the simulations (see the Materials and Methods section for details), while ERK^p(TY)^ was constructed for an active ERK with simultaneous phosphorylation of both residues. Thus, ERK^pT^, ERK^pY^, ERK^pTpY^, and ERK^pYpT^ can be regarded as transition state intermediates towards the active ERK^p(TY)^. Unphosphorylated ERKs in the absence (ERK^Apo^) and presence (ERK^Holo^) of ATP were also examined for comparison.

**Figure 1.**
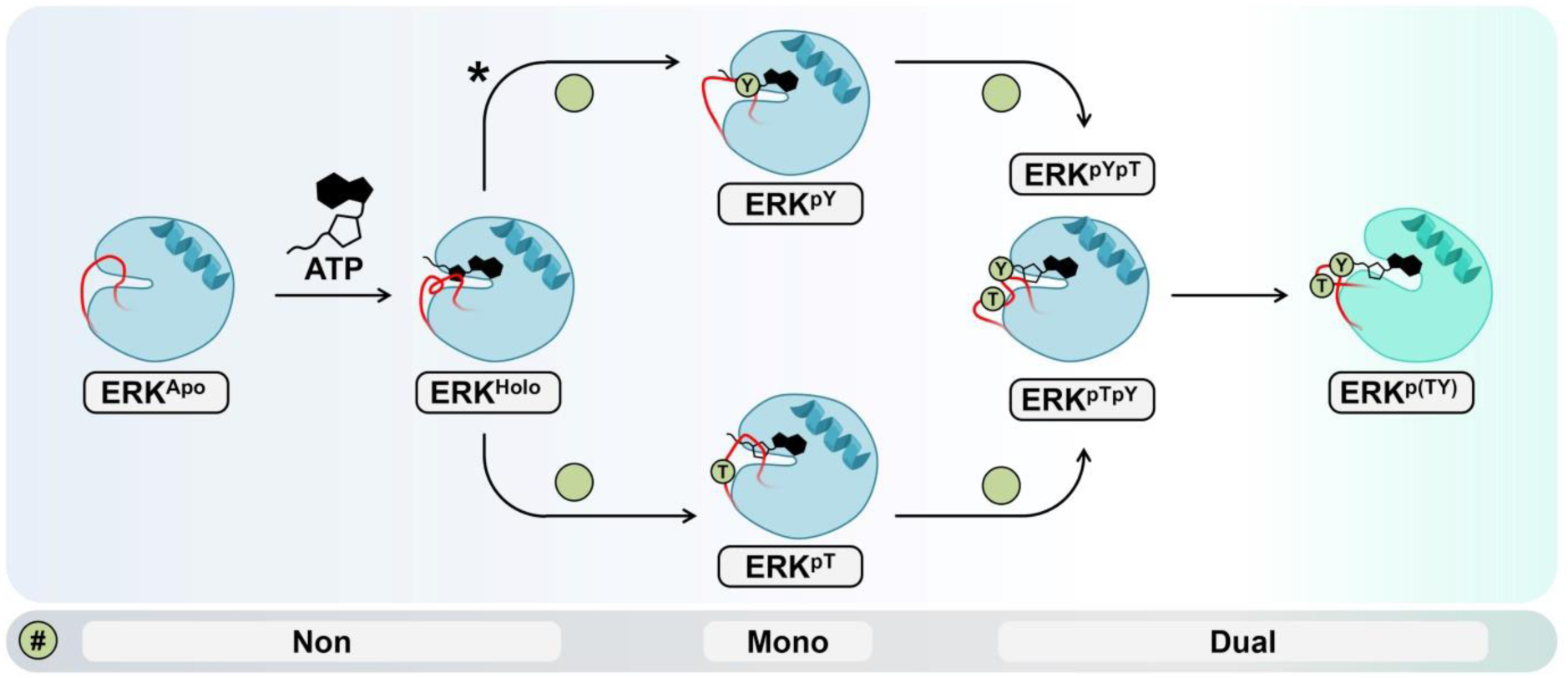
A schematic flowchart of the ERK activation mechanism. The unphosphorylated systems (ERK^Apo^ and ERK^Holo^), the monophosphorylated systems (ERK^pT^ and ERK^pY^), and the dual phosphorylated systems (ERK^pYpT^, ERK^pTpY^, and ERK^p(TY)^) were studied. All models are based on the crystal structure of phosphomimetic ERK in complex with PEA-15 (PDB ID: 4IZ5), except ERK^p(TY)^ which is based on the crystal structure of dual phosphorylated ERK (PDB ID: 6OPG). ATP is shown in black, the activation loop in red, and the phosphate group substituent as a green circle with the respective phosphorylation site noted inside. System names are given under each representation. The symbol, *, indicates the preferred pathway identified in vitro.

To evaluate the lobe dynamics and the compactness of ERK, we calculated the distance between the centers of mass (COM) of the lobes, the angle of these COMs with the hinge, and the overall height of ERK, excluding the disordered N-termini (i.e., β1-β2, residues 1-20) (**Figure 2A**). An additional energy calculation was performed to further assess the interaction between the lobes. For ERK^Apo^, as expected, the removal of ATP tends to cause expansion and dissociation of the two lobes, suggesting that ATP is the driving force to maintain the lobes in proximity, as in ERK^Holo^. The presence of ATP in the cleft assembles the C-spine and forces the lobes to close. The alignment of R-spine residues in the inactive and active states of ERK, as given by Ramachandran plots of backbone dihedral angles, suggests that compactness relies solely on the C-spine (**Figure S2**). This may imply that ERK’s transition to a more compact structure requires less conformational change compared to kinases with unaligned regulatory spine. Further changes in the compactness of the lobes were observed for the phosphorylated ERK systems, as the phosphate groups can introduce new interactions with nearby basic residues (**Figure 2B**). ERK^pY^ shows stronger interactions between the lobes, as evidenced by the decrease in interaction energy value due to the predominant binding of pY187 to R67, but the long sidechain of pY187 involved in the interaction causes slight dissociation of the lobes. In contrast, for ERK^pT^, pT185 is mostly stabilized by interacting with R191 and R194 in the C-lobe. Hence, monophosphorylation of T185 induces minor dimensional changes in the lobes, despite promoting interactions between them. Rationally, in monophosphorylated ERK^pT^ and ERK^pY^, there are fewer electrostatic interactions compared to dual phosphorylated ERK^p(TY)^. The introduction of a second phosphate group can increase electrostatic interactions, thereby enhancing the constraints on the activation loop. Indeed, due to the incorporation of an additional phosphate group as a source of electrostatic interactions, the dual phosphorylated ERK^p(TY)^ has the most compact structure and strong interaction between the lobes, consistent with the expectations for an active conformation.

**Figure 2.**
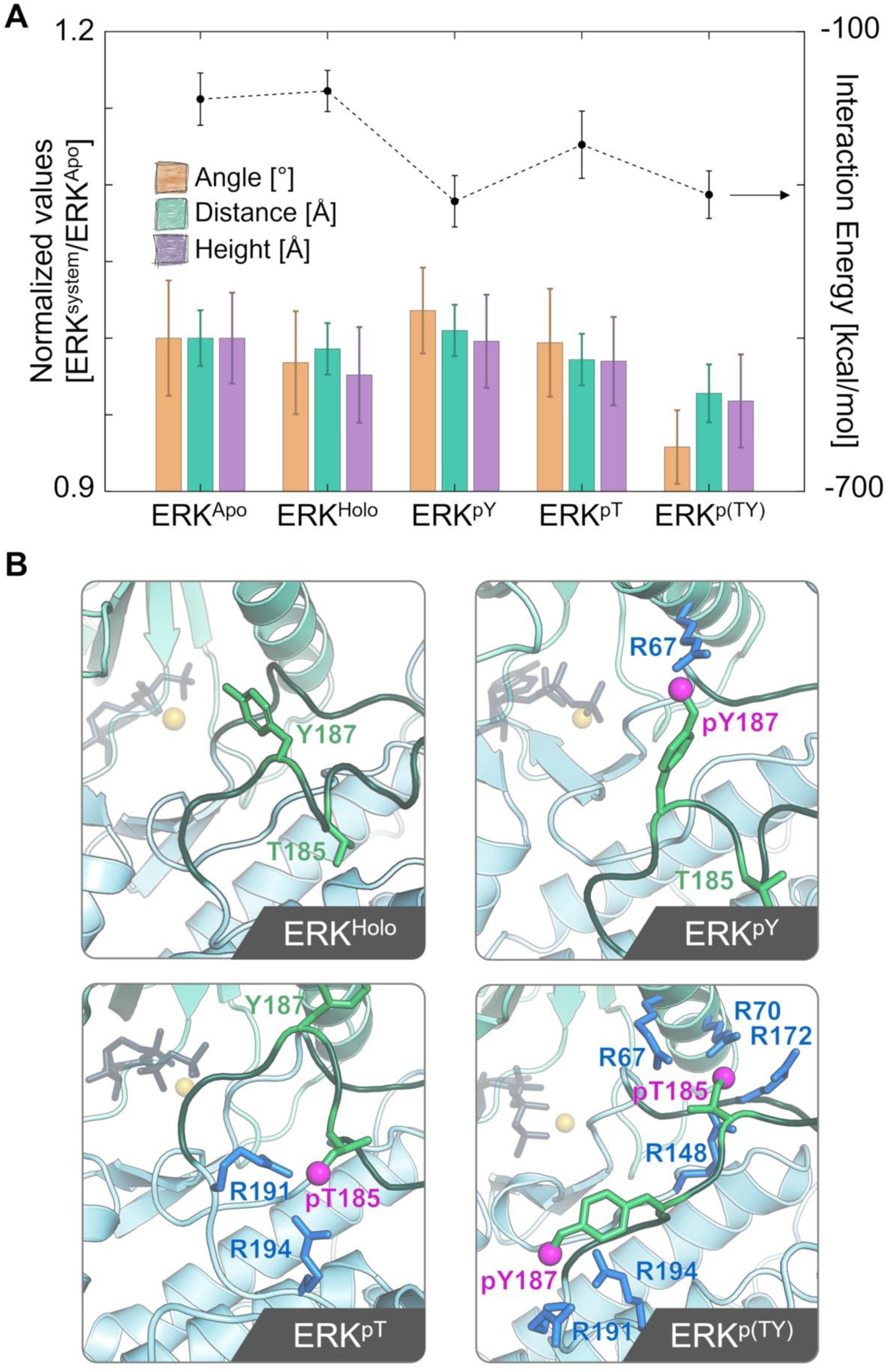
ERK compactness is stabilized by inter-lobe salt bridges. (A) Quantification of the compactness of the lobes based on three structural parameters with the scale shown on the left axis, including the distance between the center of mass (COM) of the lobes (turquoise bar), the angle formed by the COMs of the lobes and the hinge region (orange bar), and the maximum ERK height (purple bar). All values were normalized to those obtained for ERK^Apo^. A dashed line representing the interaction energy between the lobes with the scale shown on the right axis. (B) Snapshots illustrating salt bridge interactions between phosphorylated residues in the activation loop and nearby basic residues for ERK^Holo^, ERK^pY^, ERK^pT^, and ERK^p(TY)^. Small beads at the tip of the sidechain of pT185 and pY187 indicate phosphate groups.

### The ensembles of the activation loop conformations and its stability

To investigate how phosphorylation affects the conformations and dynamics of ERK’s activation loop, we performed cluster analysis to sample the ensembles of the conformation of the activation loop. For ERK^Apo^, the highly populated loop conformation (cluster #1) has an occurrence of 11%, while for ERK^Holo^, the occurrence for the highly populated loop conformation increases to 24% (**Figure 3A and Table S2**). This increase in occurrence suggests that the loop becomes more restricted to a narrower range of conformations. This narrowing of the conformational distribution is further emphasized by the cumulative occurrence plots: in ERK^Apo^, 50% of the loop conformations are grouped into 7 clusters, whereas in ERK^Holo^, it takes just 5 clusters to reach the same threshold. This trend becomes more apparent in the fully active ERK^p(TY)^, where only 2 clusters are needed to represent 50% of the loop conformational ensemble. This suggests that a phosphate group constrains the activation loop, leading to a narrower conformational distribution [46]. Monophosphorylated ERKs (ERK^pY^ and ERK^pT^) show similar trends, with a narrower distribution compared to the unphosphorylated ERK^Holo^, but a broader distribution than the dual phosphorylated ERK^p(TY)^. However, sequential dual phosphorylation shows conflicting results, both different from ERK^p(TY)^. ERK^pYpT^ shows a slight narrowing of its conformational distribution compared to its precursor ERK^pY^, but ERK^pTpY^ shows a significant broadening of its distribution compared to all phosphorylated systems. These findings highlight two important points: (i) For the experimentally determined ERK^p(TY)^, we can assume that the dynamic interaction with MEK may influence the final loop conformation. Thus, although our successive dual phosphorylation systems, ERK^pYpT^ and ERK^pTpY^ can be considered “active”, their different cluster distributions compared to ERK^p(TY)^ suggest that they may represent different active conformations within the ensemble of possible active states, and that their activation loop is expected to present a different conformation than that of the experimentally based ERK^p(TY)^. (ii) These results demonstrate that dual phosphorylation does not always restrict the activation loop to specific conformations but can also cause destabilization of certain sections of the loop, leading to a wider range of conformational possibilities. We further evaluated the conformational constraints on the activation loop by examining the geometric properties of the loop in terms of its area and elongation. On average, all systems present similar values for the loop area (**Figures 3B and S3**). However, the phosphorylated systems (both mono and dual) display narrower area distributions compared to the ERK^Apo^ and ERK^Holo^ systems. The elongation values show changes in expansion and compaction of the loop *depending on the degree of phosphorylation*. Both types of monophosphorylation force the loop to adopt a slightly more elongated conformation than observed for ERK^Apo^ or ERK^Holo^, indicating that the loop tends to expand under these conditions. Conversely, all sequentially phosphorylated systems exhibit a decrease in elongation compared to their mono-counterparts, including the ERK^p(TY)^. This suggests that dual phosphorylation forces the loop to reassume a compact conformation. This trend was previously demonstrated in our geometric measurements of lobes, suggesting that monophosphorylation may promote opening of the lobes and less compaction, while dual phosphorylation is more likely to facilitate the closing of the lobes.

**Figure 3.**
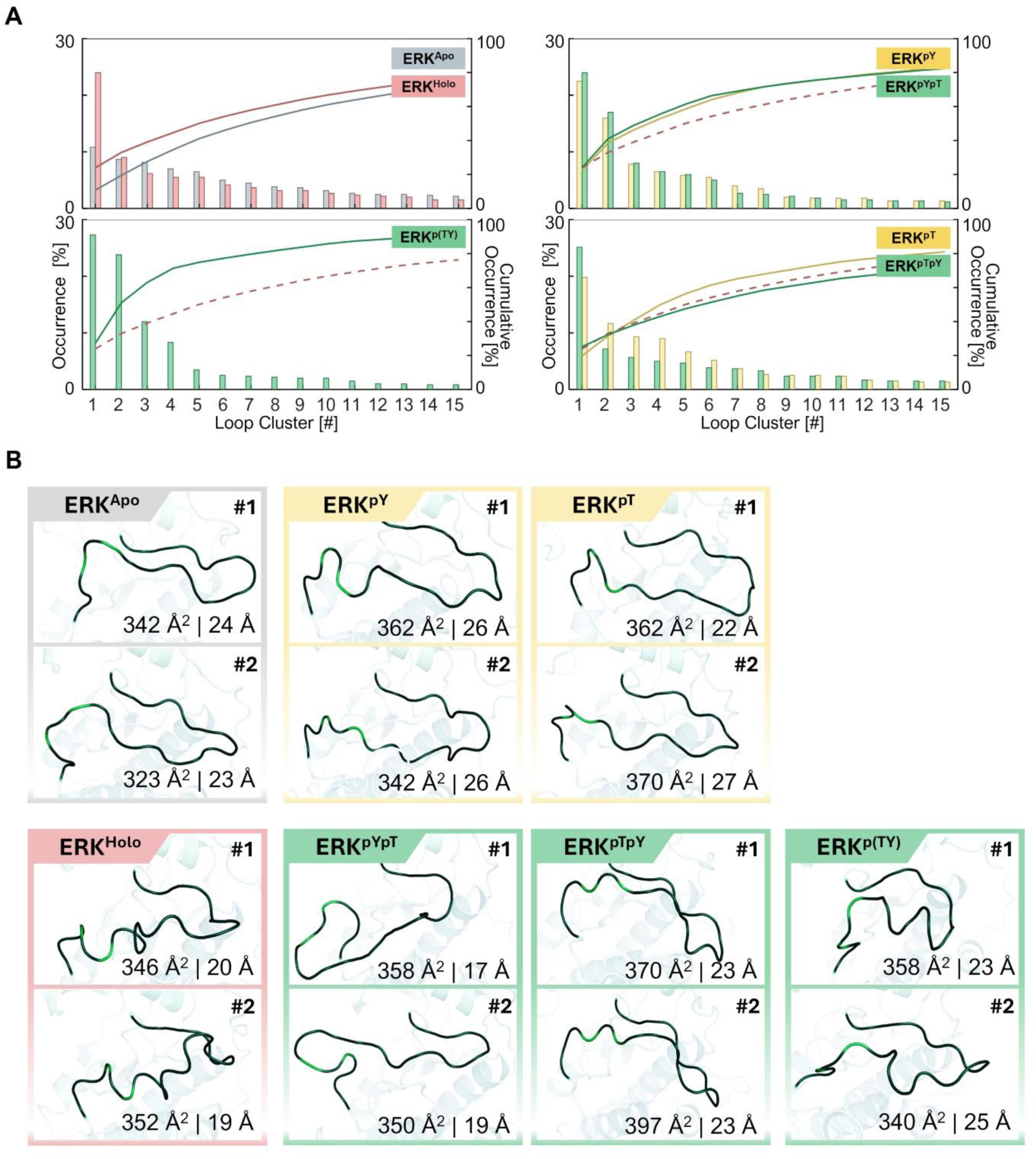
Mono and dual phosphorylation induce opposing allosteric elongations in the activation loop. (A) Bar charts showing cluster distribution by occurrence, with cumulative occurrences shown as solid lines. For comparison, the dashed red line shows the cumulative occurrences of ERK^Holo^ (inactive in the presence of ATP) clusters. (B) Representative snapshots of the activation loop for the first two clusters (#1 and #2) for the inactive systems in the absence and presence of ATP (ERK^Apo^ and ERK^Holo^), the monophosphorylated systems (ERK^pT^ and ERK^pY^), and the dual phosphorylated systems (ERK^pYpT^, ERK^pTpY^, and ERK^p(TY)^). The calculated values for loop area and elongation are marked in each snapshot.

We calculated the root mean square fluctuation (RMSF) of the activation loop to evaluate how different phosphorylation codes influence the loop stability in terms of fluctuations. Since the activation loop is composed predominantly of uncharged residues (about 75%), its stabilization relies on hydrogen bonding and weak hydrophobic interactions, increasing its tendency for frequent conformational changes. As expected, the loop in ERK^Apo^ shows a typical semi-Gaussian curve, where the centered residues of the activation loop (residues 178-185) present higher fluctuation values compared to the edges (residues 167 and 190) (**Figure 4A**). Noticeably, the activation loop in ERK^Holo^ shows less fluctuations, indicating a more stable loop compared to ERK^Apo^. This stabilization results from closing the lobes in response to ATP insertion (**Figure 2**). ERK^p(TY)^ exhibits an even greater stabilization of the loop, with a greater reduction in its fluctuations. The closed structure constrains the loop to occupy a smaller space, resulting in a shorter elongated conformation (**Figure 3 and S3**). This is not the case for monophosphorylation, where ERK^pT^ presents similar stability to ERK^p(TY)^, but ERK^pY^ unexpectedly destabilized the loop. Interestingly, after successive phosphorylation (of ERK^pYpT^ and ERK^pTpY^), the different curves of RMSF for ERK^pY^ and ERK^pT^ merge into a single fluctuation curve that resembles the stable curve of the active ERK^p(TY)^. The greatest reduction in fluctuation for all the curves is observed at the phosphorylation sites (T185 and Y187), confirming that stabilization of the flexible loop, an allosteric effect, can be achieved by the addition of phosphate groups. The unphosphorylated residues, T185 and Y187, have weak interactions, but upon phosphorylation, the interactions are strengthened for both lobes, specifically in the electrostatic component (**Figure 4B**). Post successive phosphorylation reveals different trends for the initially monophosphorylated residues. In ERK^pTpY^, the latter phosphorylation on Y187 strengthens the interactions of both phosphorylated residues with the lobes. In ERK^pYpT^, the latter phosphorylation on T185 strengthens its interaction but also decreases the interaction energy of pY187 with the lobes. This suggests that pY187 has weaker interactions compared to those of pT185, with the two residues competing for binding the same residues.

**Figure 4.**
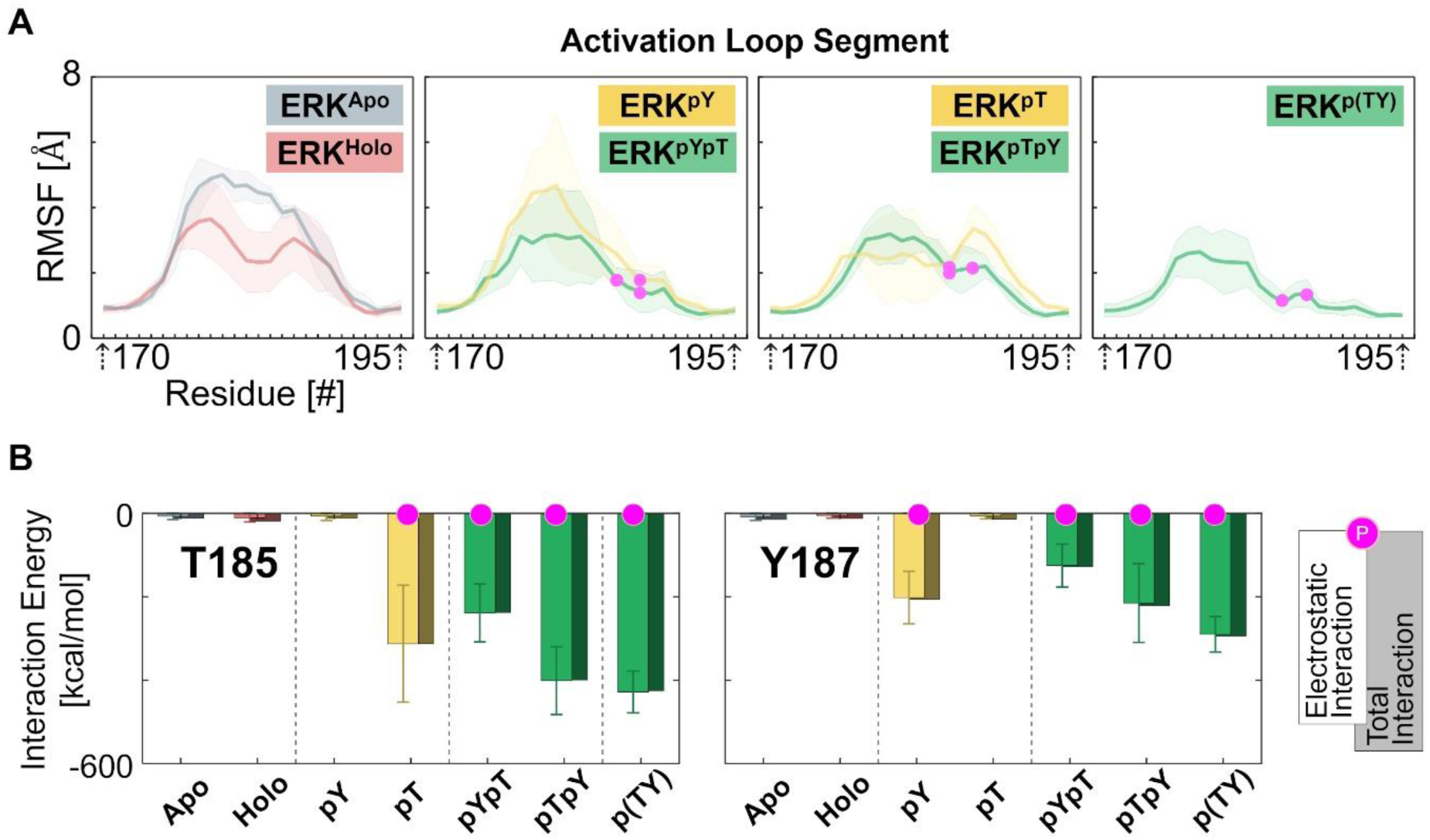
Enhanced stability of the activation loop through the introduction of phosphate groups. (A) Root mean square fluctuation (RMSF) of the activation loop, and (B) total interaction energy of the phosphorylation sites with the kinase, highlighting the electrostatic interaction, for the inactive systems in the absence and presence of ATP (ERK^Apo^ and ERK^Holo^), the monophosphorylated systems (ERK^pT^ and ERK^pY^), and the dual phosphorylated systems (ERK^pYpT^, ERK^pTpY^, and ERK^p(TY)^). In the RMSF plots, the average value per residue is plotted as a thick solid line and the standard deviation is illustrated as its faded background color. Phosphorylated residues are shown as magenta circles.

### Phosphorylation induces allosteric effects

The observation that ERK^pT^ has a similar RMSF curve to that of ERK^p(TY)^, raises an intriguing question. If the stabilities are similar, why does ERK^pT^ need further phosphorylation at Y187 to become active? To address this question, we focused on the flexible regions, including the hinge and L16 segments, to observe how these regions are affected by the different phosphorylation codes in the activation loop. We reasoned that the fluctuating behavior of the activation loop can propagate through the kinase domain, and thus these regions should be sensitive enough to show changes in their fluctuations. Our findings confirmed that allosteric effects promoted by the loop echo through all segments and affect the stability of these regions (**Figure 5A**). A thorough examination of the RMSF curves of the hinge and L16 segments reveals that ATP insertion, which triggered the allosteric effect, leads to a decrease in fluctuation in the hinge and an increase in L16. As ATP induces lobe closure, the hinge rigidifies and exhibits a reduction in fluctuations. Monophosphorylated ERK^pT^ and ERK^pY^ show no change in the fluctuations of the L16 segment compared to ERK^Holo^, but the hinge segment exhibits destabilization, which can be attributed to the opening of the lobes as seen above. However, unlike the triggered loop fluctuations (**Figure 4A**), where each phosphorylation site resulted in a different curve, monophosphorylated ERK^pT^ and ERK^pY^ show similar averaged fluctuation patterns per region for the hinge and L16 segments. All dual phosphorylated ERKs also show similar curves per region with less fluctuations compared to the monophosphorylated systems. The regional fluctuation curves successfully demonstrate that phosphorylation of the activation loop constrains the kinase domain structure, affecting the behavior and stability of distant domains. To further assess the impact of the phosphorylation code, we mapped the spatial proximity of residue pairs between the activation loop and the rest of the kinase domain (**Figure S4**). We identified two regions with notable changes in their contacts, the region containing HRD (residues 144-154) and the P+1 segment (residues 197-207). Contact maps revealed that in non-dual phosphorylated systems, ERK^Holo^, ERK^pT^ and ERK^pY^, HRD residues 148-149 contact the activation loop (residues 182-184), but dual phosphorylated ERK^pTpY^, ERK^pYpT^, and ERK^p(TY)^ lack these contacts (**Figure 5B**). These differences clarify the significance of dual phosphorylation in the activation mechanism of ERK. In ERK^Holo^, ATP induces lobes closure, positioning the HRD segment in proximity to the activation loop. This event is also supported by significant reduction in fluctuations of residues 182-184 as observed in the fluctuation curve (**Figure 4A**). Proximity to the HRD is also present in monophosphorylated ERKs, where the loop adopts a more elongated conformation. However, after adopting a compact conformation due to dual phosphorylation, the loop detaches from the HRD segment, allowing its residues to readjust and regain a stable conformation (**Figure S5**). The P+1 segment is located beneath the activation loop, and its contacts are expected to change depending on the loop conformation. Both ERK^Holo^ and ERK^p(TY)^ have similar contacts between the P+1 segment and the activation loop (**Figure 5C**). ERK^pT^ has a slight shift in contacts while maintaining an overall interaction pattern similar to ERK^p(TY)^, but these contacts are weak in ERK^pY^. For dual phosphorylated ERKs, ERK^pYpT^ has a similar contact pattern to those of ERK^Holo^ and ERK^p(TY)^, while ERK^pTpY^ loses these contacts, although its precursor ERK^pT^ retains them. This discrepancy suggests that the order of successive phosphorylation events may have different impacts on regions that contact the activation loop, leading us to investigate whether there is a preferred sequential phosphorylation pathway for ERK activation.

**Figure 5.**
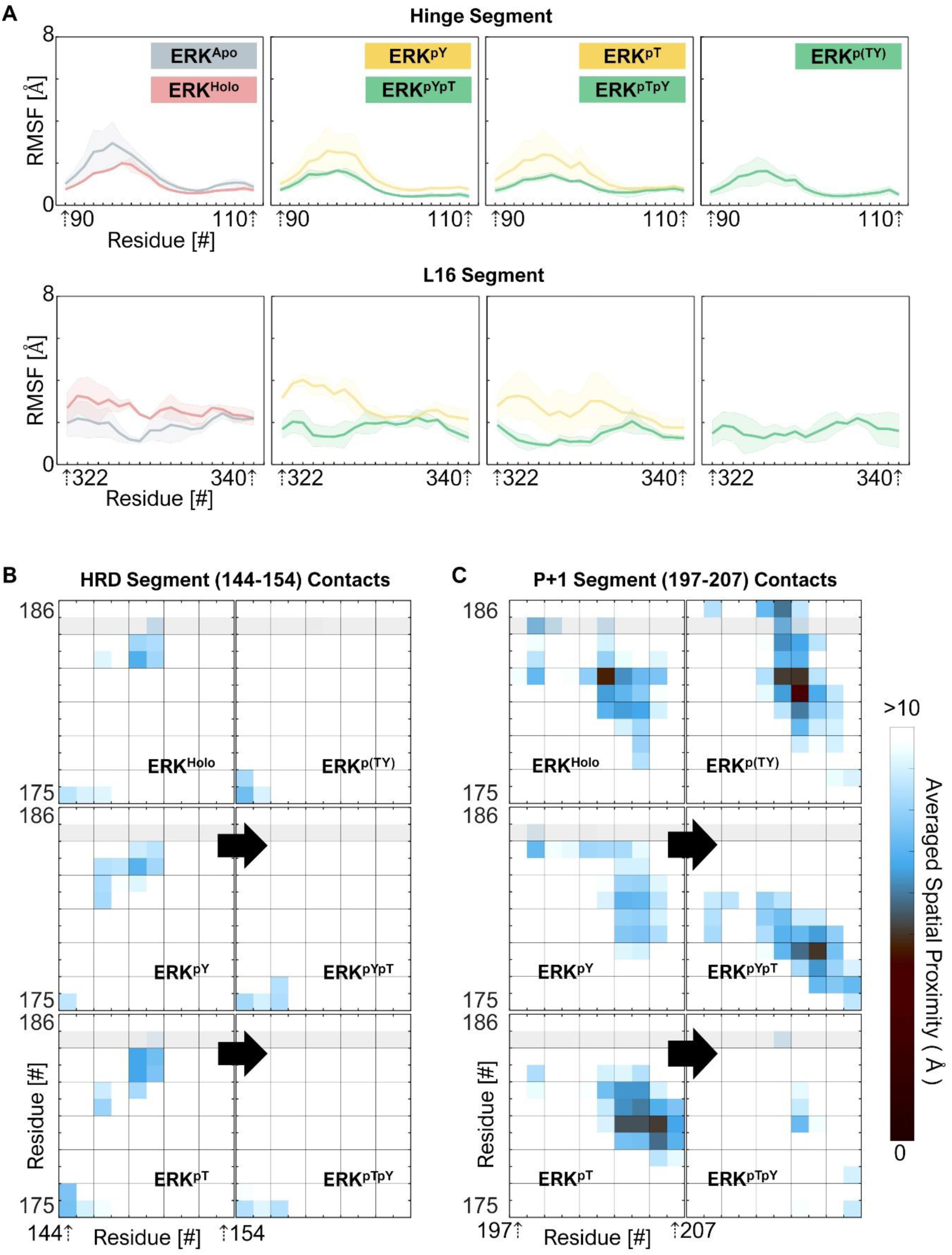
Phosphorylation code triggers allosteric effects, illustrating how activation affects the stability of binding and catalytic sites. (A) Root mean square fluctuation (RMSF) of two flexible regions, the hinge and L16 segments, for the inactive systems in the absence and presence of ATP (ERK^Apo^ and ERK^Holo^), the monophosphorylated systems (ERK^pT^ and ERK^pY^), and the dual phosphorylated systems (ERK^pYpT^, ERK^pTpY^, and ERK^p(TY)^). The average value per residue is plotted as a thick solid line and the standard deviation is illustrated as its faded background color. Contact maps for the activation loop versus (B) the region containing HRD (residues 144-154) and (C) the P+1 segment (residues 197-207).

### Accessibility of the monophosphorylation sites

Numerous studies identified Y187 as the predominant first phosphorylation site [21–23, 47–49]. They either found massive accumulation of pY187 or identified pT185 intermediates, not by a direct monophosphorylation of ERK, but as a result of dephosphorylation of dual phosphorylated ERK. This opens the possibility of selective phosphorylation in the activation loop of ERK. To understand why monophosphorylation occurs predominantly at Y187, we calculated the orientation of the sidechain of the phosphorylated residue and its solvent access surface area (SASA) to evaluate how well each phosphorylation site can interact with its environment including the surrounding solvent. In ERK^Apo^, T185 and Y187 have similar average orientation values of ∼100° (**Figures 6A and S6**). Upon ATP binding, the lobes adopt a more closed structure, subjecting the activation loop to steric constraints and conformations in which the two residues have different orientation angles. In ERK^Holo^, while T185 faces the kinase domain, Y187 is predominantly exposed and faces the solvent. For the pT185 → pY187 phosphorylation pathway, pT185 and Y187 in ERK^pT^ retain orientations resembling those observed in ERK^Holo^. Upon cascading Y187 phosphorylation, there is a significant change in the orientation of pT185 in ERK^pTpY^, with lower orientation angles indicating that it faces the kinase domain in a more hindered orientation. This angle reduction is not observed in ERK^p(TY)^, confirming that the dynamic interaction with MEK is responsible for the conformation of the activation loop of active ERK. In contrast, the pY187 → pT185 phosphorylation pathway yields different outcomes. In ERK^pY^, the initial phosphorylation at Y187 significantly affects the orientation of T185, positioning it exclusively toward the solvent. This change makes T185 more solvent accessible, thus easier to phosphorylate in the subsequent phosphorylation step. The change in T185 orientation correlates with an increase in SASA (**Figure 6B**). Our results provide a rational explanation for ERK’s selective activation likely follows *pY187 → pT185 phosphorylation*, consistent with experimental results [22, 23]. This can be explained by two observations (**Figure 6C**): (i) Y187 is more solvent accessible than T185, making it more likely to be phosphorylated first, and (ii) by following this pathway, the hindered T185 can be exposed and more easily phosphorylated.

**Figure 6.**
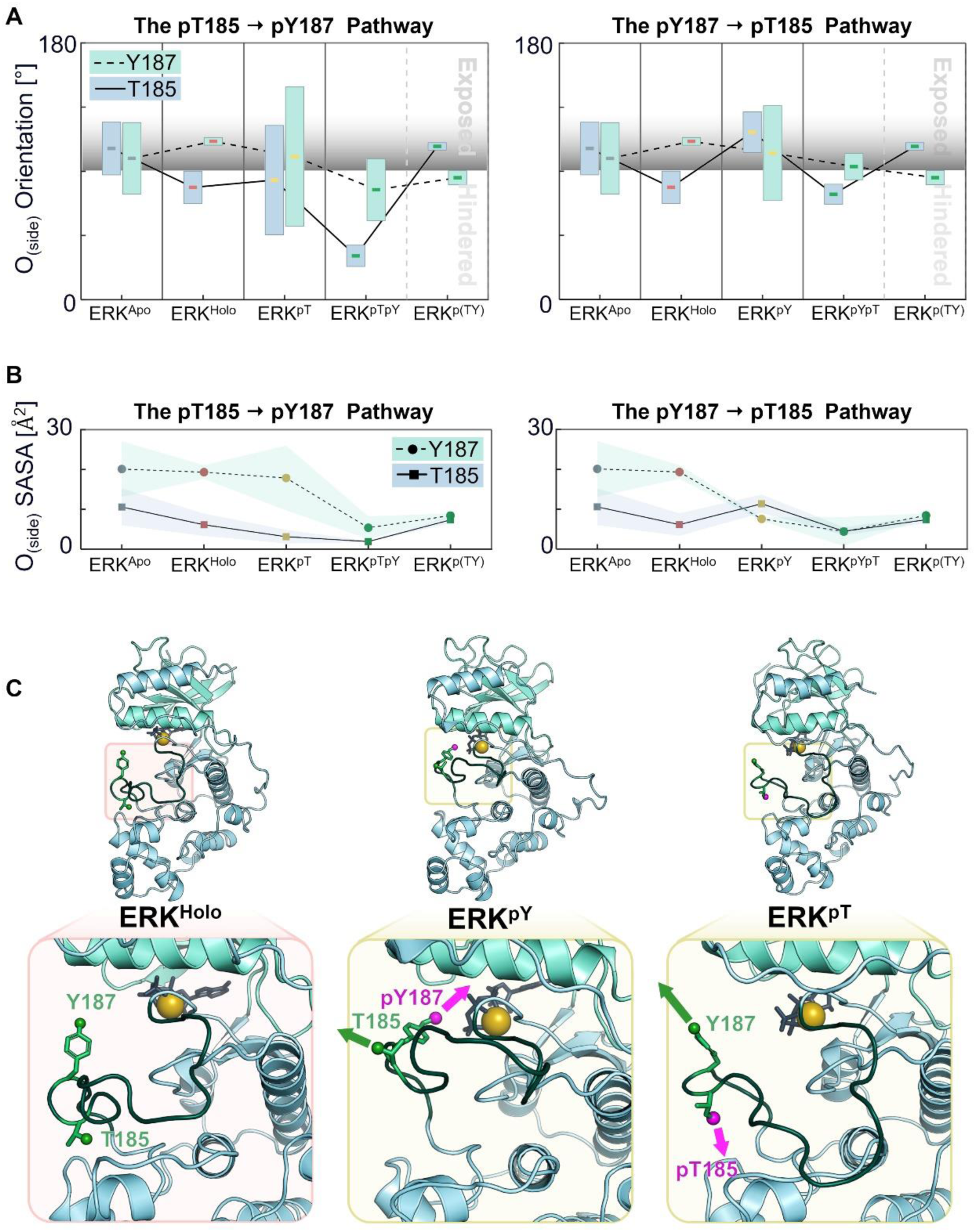
Accessibility of phosphorylation sites favors the pY187 → pT185 pathway. (A) The orientation angles of the sidechains of T185 (blue, solid line) and Y187 (teal, dashed line) with respect to the αF-helix for different phosphorylation pathways. Values above 90° indicate that the atom is facing the solvent. (B) Averaged solvent access surface area (SASA) of the side group oxygens of T185 (solid line) and Y187 (dashed line) for different phosphorylation pathways. The standard deviation is illustrated as its faded background color. (C) Snapshots of the phosphorylation sites for ERK^Holo^, ERK^pY^, and ERK^pT^. The phosphorylated residue is shown as a magenta bead.

### The successive phosphorylation pathways affect the activation site’s binding conformation

ERK phosphorylation by MEK is currently classified as random, implying that MEK is capable of phosphorylating T185 and Y187 as a first step [21, 49]. Theoretically, these events have an equal chance of occurring. However, we found that the initial phosphorylation of ERK is selective, with Y187 preferentially phosphorylated first, making the pY187 → pT185 phosphorylation pathway more likely. This finding raised an essential question: if Y187 is the preferred initial phosphorylation site, does its subsequent phosphorylation on T185 activate ERK more effectively than the alternative pathway, pT185 → pY187? To address this question, we focused on the activation site to identify the specific roles of these residues and how they affect ERK activity. We mapped the hydrogen bonds and electrostatic interactions that bind ATP in its pocket. Of all the possible atomic pairings based on proximity, only a small number of residues show consistency in their binding occurrence across all replicates in the set (**Table S3**). The distribution curves represent the distances for each pairing, and the distribution patterns are compared to those of inactive ERK^Holo^ and active ERK^p(TY)^. For example, two hinge residues, Q105 and D106 stabilize ATP through hydrogen bonding with the adenosine’s amine (**Figure S7**). These residues exhibit similar interaction distribution curves across all systems, suggesting that they are unaffected by the progression of the activation process. It is important to note that although D106 binds ATP using its backbone carbonyl, its sidechain was previously found to impact ERK activity as well [50, 51]. Another residue, R67 which is located outside the activation site has been shown to interact with ATP and the phosphorylation sites (**Figures 2 and S7**). Its interactions with the phosphorylation sites depend on the type of monophosphorylation. For Y187 as the initial phosphorylation site, R67 exhibits binding patterns like the active state, whereas for T185, it produces patterns that fit the inactive state. Based on its distinct distributions, we suggest this residue as an experimental flag for the identification of the initial phosphorylated site. The pairing of K151 with O^γ^ of ATP in the inactive conformation ERK^Holo^ yields a distribution consisting of three populations separated by 3.8 and 5.9 Å thresholds (**Figure 7A**). The active conformation, however, has only a single population below the 3.8 Å threshold. These differences indicate that in the inactive state, the interaction of K151 with ATP is expected to be weaker than in the active state due to the larger distance population in ERK^Holo^ compared to ERK^p(TY)^. In monophosphorylated ERKs, the interaction of K151 with ATP is not affected by the phosphorylation at Y187 or T185, as shown by the high distribution values of ERK^pY^ and ERK^pT^. This is consistent with the contact maps for the HRD segment, in which K151 changes its contacts only in the dual phosphorylated ERKs when the segment and loop are dissociated. After successive phosphorylation, the distance distributions revealed that K151 adopts a single distance population, but with the average distance shifting to larger or shorter distances: in ERK^pYpT^, the population correlates with the active state and presents stronger interactions at shorter distances, while ERK^pTpY^’s larger distance population is more consistent with the inactive state. The distance populations show that K54 near the G-loop binds ATP strongly in the inactive state, but this interaction weakens in the active state. The distribution patterns show that monophosphorylated ERK^pY^ resembles the inactive state, while ERK^pT^ resembles ERK^p(TY)^. Remarkably, successive phosphorylation alternates this trend, with K54 loosely binding ATP in ERK^pYpT^ and *vice versa* for ERK^pTpY^. This is reflected in the K54-E71 distance distribution, where this weak salt bridge in ERK^pY^ is strengthen in ERK^pYpT^, in contrast to ERK^pY^ and ERK^pTpY^. Thus, when ERK is dual phosphorylated, its conformation appears confined to a constant equilibrium between two major ensembles (**Figure 7B**). The ensembles include a less active conformation in which K54 does not bind E71 and there is minimal interaction between the active site and the HRD segment (the pT185 → pY187 pathway), and a more active conformation in which ATP is locked in its pocket by the K54-E71 salt bridge, and the activation site interacts with the HRD segment (the pY187 → pT185 pathway).

**Figure 7.**
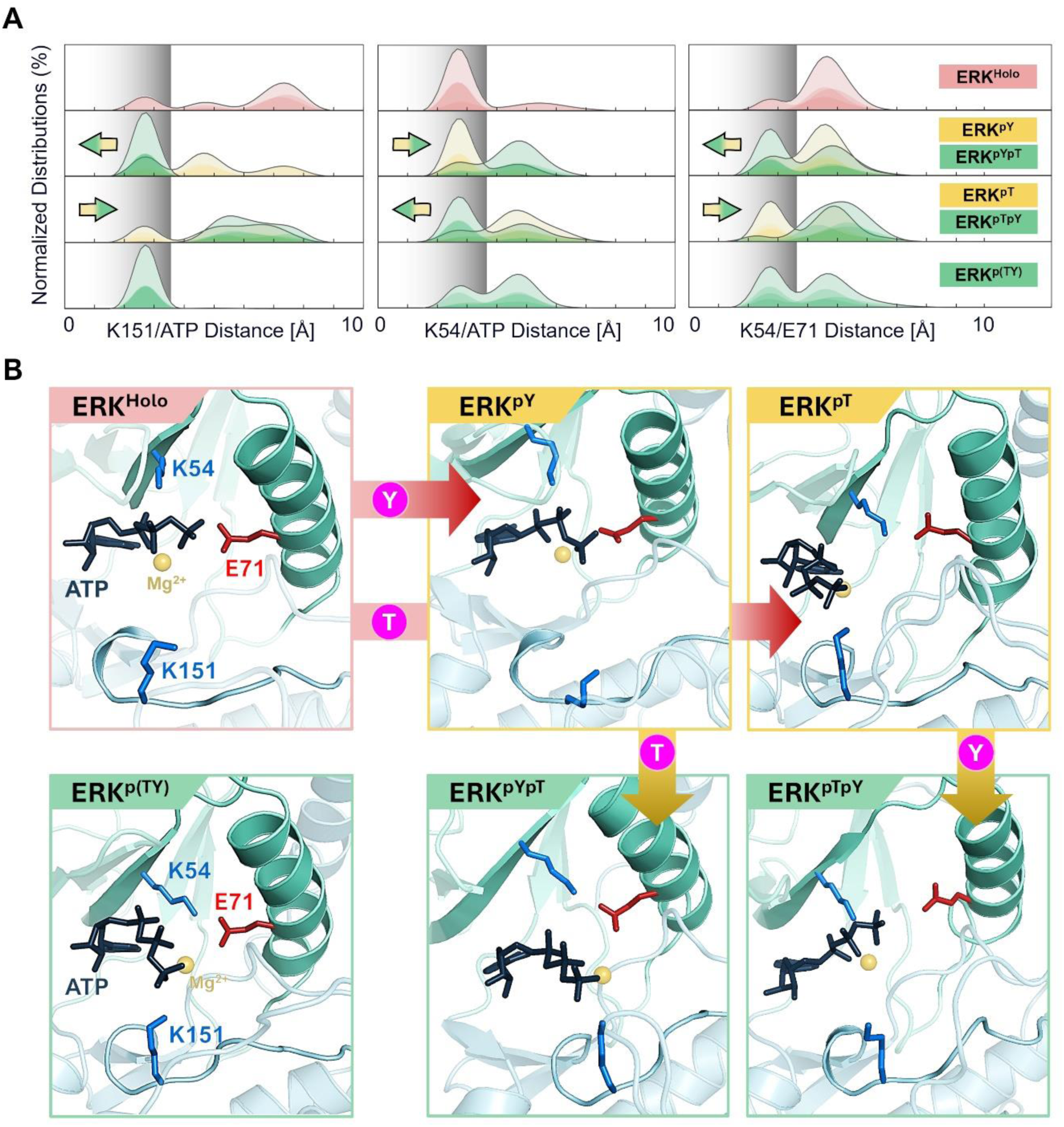
The pY187 → pT185 pathway derives the binding pattern of an active state. (A) Distributions of normalized distances from ATP to K151 and K54, and from K54 to E71. The distributions show that the distances can be divided into two types of populations based on their similarity to the inactive state (ERK^Holo^) and the active (ERK^p(TY)^). Arrows indicate the direction of population shift. (B) Snapshots of the ATP binding site illustrating the formation of the K54-E71 and ATP-K151 salt bridges for the inactive system in the presence of ATP (ERK^Holo^), the monophosphorylated systems (ERK^pT^ and ERK^pY^), and the dual phosphorylated systems (ERK^pYpT^, ERK^pTpY^, and ERK^p(TY)^).

## Discussion

ERK is among the most diverse kinases with hundreds of direct substrates, including transcription factors and signaling proteins, making it a central kinase in the cell, at the heart of cell proliferation, and a key regulator in multiple pathways. ERK mutations are rare in cancer, likely due to OIS (oncogene induced senescence). Its role is crucial in cancer and in other pathologies, including (neuro)developmental disorders, making the understanding of its regulation vital [52]. While a few ATP-competitive drugs (e.g., LY3214996, Ulixertinib (BVD-523), and Ravoxertinib (GDC-0994)) have been, or are, in clinical trials, unlike MEK, no direct allosteric drugs have been developed for ERK. Currently, it is targeted indirectly, via MEK, as well as via e.g., ErbB receptors and PI3K/mTOR pathway inhibitors. Targeting ERK is challenging because of the difficulty in developing highly selective inhibitors thus expected off-target effects, and the emergence of drug-resistant mutations [53].

Our aim is gaining in-depth understanding of its behavior. Deciphering the phosphorylation signaling code of ERK at the fundamental conformational behavior level is our goal, with the hope that it will help in drug development [54, 55]. Recently, our mechanistic studies of mTOR, and SHP2 yielded some pharmacological cues [30, 56]. Here, we screen modeled ERK components during activation: unphosphorylated ERK in absence and presence of ATP, two monophosphorylated ERKs, two successive phosphorylated systems, and an experimental reference active system.

### Dual phosphorylation enables the catalytic segment

Here we identified two simple, yet critical requirements for ERK to function efficiently: (i) a compact activation loop structure that minimizes steric hindrance and enhances substrate accessibility to the catalytic site [45], and (ii) unimpeded interaction between the catalytic HRD motif and ATP in the binding pocket. In the presence of ATP, the activation loop of ERK adopts a compact conformation restricted and confined by the lobes closure, facilitating its interaction with substrates (**Figure 2**). The compact structure also enables the interaction with its upstream activator MEK to initiate the activation process. However, ATP presence in the active site interferes with ERK catalysis by destabilizing the HRD segment, pushing it towards the activation loop (**Figure 5**), making the coexistence of the two features seems unlikely. To resolve this conflict, ERK undergoes a two-step phosphorylation process. The initial phosphorylation step introduces new constraints that limit the activation loop dynamics causing it to disfavor the compact state toward a more elongated conformation (**Figure 3**) [22]. We believe that the destabilization and elongation of the loop at this step is a rational mechanistic step designed to overcome the constraints and to increase the likelihood of successive phosphorylation. Dual phosphorylation saturates the loop with salt bridges that force it to recompact without interfering with the HRD motif. This final configuration restores the ability of the HRD segment to interact effectively with the activation pocket while keeping the catalytic site accessible for the substrates (**Figure 8**).

**Figure 8.**
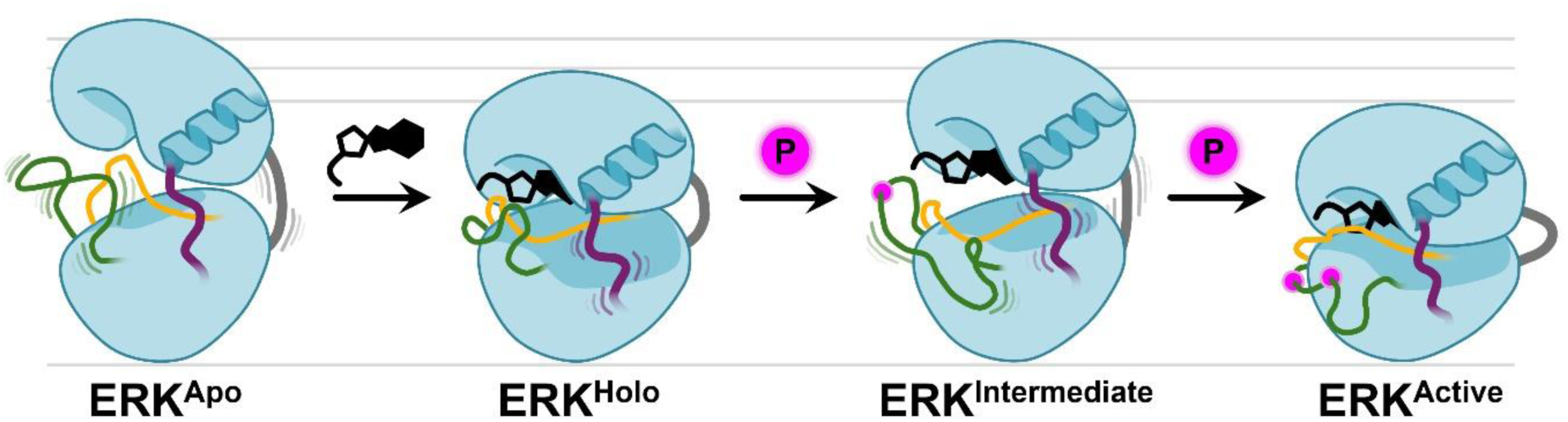
Schematic illustration of the activation mechanism. Dual phosphorylation recompacts the activation loop, allowing HRD to interact with ATP. The activation loop is shown in green, the L16 segment in purple, the hinge in gray, and the region containing HRD in orange. Changes in kinase dimensions are emphasized by a gray horizontal ruler.

### Phosphorylation allosterically orchestrates distant regions to favor dual phosphorylation

ERK possesses two flexible regions on top of its activation loop. These regions are the hinge segment connecting the N-lobe and C-lobe which facilitates transitions between the open and closed conformations, and the L16 segment which is the binding site for homodimerization of ERK [14, 50]. Studies have demonstrated that these flexible regions are susceptible to allosteric effects that allow the kinase to effectively respond to changes in its cellular environment [19]. We demonstrated that the impact of the phosphorylation steps can propagate throughout the kinase, affecting these flexible regions allosterically, thereby regulating activation (**Figure 8)**. *Importantly, the behavior of these distant segments is not dictated by the phosphorylation order, but rather by a tri-state signaling code based solely on phosphorylation degree of the activation loop: unphosphorylated, monophosphorylated, and dual phosphorylated* (**Figure 5**). Upon ATP binding, the kinase becomes committed to catalysis [57]. At this step, the hinge is stabilized, minimizing lobe dissociation and ensuring ATP binding. At the same time, the homodimerization site becomes unstable, potentially minimizing homodimerization and facilitating interactions with hetero proteins, such as its upstream activator MEK. Following activation, the homodimerization binding site restabilizes to inspire ERK dissociation from MEK by enhancing the formation of homodimers [12, 18, 58].

### The roles of the phosphorylation sites

The ATP binding pocket serves as the most important regulatory site through which ERK fulfills its kinase role. We identified the K54-ATP salt bridge as the dominant interaction in the inactive conformation of the active site. In the active conformation there are two salt bridges, K54-E71 and K151-ATP. Mechanistically, K54 interaction with ATP weakens in favor of E71, but K151 compensates the lost interaction as an alternative stabilizer to maintain ATP aligned. The formation of the K54-E71 salt bridge stabilizes the G-loop over ATP, minimizing the chances for ATP escape. As we examined the patterns of the monophosphorylated systems, we revealed the impact of each phosphorylation site on the conformation of the activation site (**Figure 7**). *Tyrosine phosphorylation drives the conformational equilibrium toward populating the inactive binding conformation, while threonine phosphorylation drives it toward the active one*. These roles were amplified after sequential phosphorylation as monophosphorylated systems altered their binding pattern to reflect the opposite state. These roles agree with observations that were previously reported by an experimental study based on various activation loop analogues [50].

### Effective activation is order-dependent

ERK activation can follow two dual phosphorylation pathways: pY187 → pT185 (tyrosine first-threonine second) and pT185 → Y187. On the conformational level, our analysis of the activation site interaction patterns identified the first pathway as the dominant activating, while the second results in an inactive conformation (**Figures 7**). Consistent with experimental results, ERK selectively follows pY187 → pT185 [21–23, 47–49]. Our conformational analysis also provides the rational as to why it is more favorable: (i) tyrosine is more accessible than threonine (**Figure 6**), (ii) phosphorylating tyrosine first promotes a conformational change that exposes threonine, facilitating the second phosphorylation, and (iii) tyrosine is a weaker nucleophile than threonine; Hence, pY187 can easily dissociate its electrostatic interactions upon threonine’s phosphorylation to allow conformational change that stabilizes both phosphorylated sites simultaneously (**Figures 4 and S8**). The chances for threonine to change its interaction partners are lower and will require overcoming a higher energy barrier. It is reasonable to assume that nearby proteins and the formation of ERK-based dimers can control these structural constraints and eventually allow the activation loop to transition between different dual phosphorylated conformations (i.e, ERK^pTpY^, ERK^pYpT^, and ERK^p(TY)^). This translates to an equilibrium between an effectively active ensemble that can phosphorylate substrates efficiently, and a moderately active ensemble, a dormant-like-type state that can undergo conformational changes to become effectively active. This transition can be triggered by substrate binding or changes in cellular environments that allosterically promote reorganization of residues within the ATP pocket. This concept proposes an additional layer of self-regulation for ERK activity and its ability to function in various cellular environments.

To conclude, here we proposed an allosteric phosphorylation code in ERK activation. We clarified the reasons for two phosphorylation sites and their preferred phosphorylation order on the conformational level. Over a decade ago, we discussed allosteric post-translational modification (PTM) codes and their key regulatory roles [59]. Here we uncover the interplay between two phosphorylations acting through modifying the strength of the interactions of the lobes and ATP binding, stabilizing the closed, active state structure of crucial kinases in the cell, with over 600 confirmed substrates, and estimated many more [5]. This site-ordered allosteric phosphorylation PTM code is an extremely effective vehicle working efficiently by harnessing *the extent of phosphorylation to fine-tune catalysis*, to the best of our knowledge to date, an unknown mechanism. That efficiency is paramount for ERK can be gleaned from ERK1/2 being the sole target MEK. Mechanistically, this raises the possibility of targeting ERK at the tyrosine phosphorylation site by wielding protein tyrosine phosphatases (PTPs) in treatment for Ras/ERK based cancers [60]. Broadly, focusing on this site and the allosteric phosphorylation code mechanism could be considered as a potential allosteric therapeutic strategy.

## Materials and Methods

### Construction of the inactive and active ERKs

To generate the initial configurations of full-length ERK, we adopted the crystal structure of phosphomimetic ERK in complex with PEA-15 (PDB ID: 4IZ5) [61] for an inactive conformation and dual phosphorylated ERK (PDB ID: 6OPG) [42] for an active conformation. Any mutated residues present in the crystal structures were changed back to the wild-type sequence. ATP and magnesium ion (Mg^2+^) were superimposed with the existing crystalized ligands at the ATP binding-pocket. All ERK systems in this study were designed using Discovery Studio software (BIOVIA, San Diego, USA). Four different initial configurations for inactive ERK were constructed based on the crystal structure in 4IZ5, including unphosphorylated ERKs in the absence (ERK^Apo^) and presence (ERK^Holo^) of ATP, and monophosphorylated ERKs (ERK^pT^ and ERK^pY^, where T and Y denote the monophosphorylating site) in the presence of ATP. To introduce the second phosphate group to ERK^pT^ and ERK^pY^, we extracted the last structures of the monophosphorylated ERKs after the 3 µs simulation for each system and tagged them with an additional phosphate group, generating two successive phosphorylated ERKs, ERK^pTpY^ and ERK^pYpT^. To construct an active ERK, the initial coordinates were taken from the crystal structure in 6OPG, generating the active, dual phosphorylated ERK^p(TY)^. See **Figure 1** and **Table S1** for a schematic representation of the activation process of ERK and an overview of the systems for MD simulations.

### All-atom explicit MD simulations protocol

The all-atom MD simulations were performed for all systems, employing the NAMD 2.14 package [62] with the CHARMM all-atom additive force field (version C36m) [63, 64]. The simulation protocol in this work closely follows that used in our previous works [26, 31, 65–69]. The explicit TIP3 water model was used to solvate the systems within a periodic box with at least 15 Å of solvent from each edge of the kinase. The charge of all potential titratable groups was fixed at values corresponding to neutral pH. A cubic simulation box with periodic boundary conditions with the nearest image convention was used. To compute the van der Waals (vdW) interactions, the atom pair cutoff distance was set to 12.0 Å. To avoid discontinuities in the potential energy function, nonbonding energy terms were forced to slowly converge to zero, by applying a smoothing factor from a distance of 10.0 Å. Na^+^ and Cl^−^ were added to each system to neutralize the charge and to maintain physiological condition of 0.14 M.

Prior to the productions runs, a series of minimization cycles were performed on the solvents, including the ions, with a harmonically restrained protein backbone until the solvent reached 310 K. Subsequent preequilibrium simulations were performed with dynamic cycles while gradually releasing the harmonic restraints on the protein backbones. The interaction potentials between the atoms were calculated by the short-range vdW interactions using switching functions. For the long-range electrostatic interaction, the particle mesh Ewald (PME) method with a grading space of 1.0 Å was used. In the production runs, the constant temperature of 310 K was maintained by the Langevin thermostat method with a damping coefficient of 1 ps^−1^ and the pressure at 1 atm was sustained by the Nosé-Hoover Langevin piston pressure control algorithm. Covalent bonds including hydrogen atoms were constrained using the SHAKE algorithm. MD simulations were performed for each system under the NPT ensemble (constant number of atoms, pressure, and temperature) with 2 fs time step. Each system was subjected to three independent simulations, each running for 3 µs, with trajectories recorded at 1 ns intervals. Systems were verified for convergence and reaching a stable conformation for the kinase domain by Cαs RMSD values (±1 Å).

### Software usage and analyses

All analyses focused on the last 2 µs in which the systems were converged. Systems were analyzed independently for each run, resulting in 2000 data points per replicate and 6000 data points per system (each system was run three times). The results are presented in two formats: (i) mean values ± standard deviations, calculated based on the three averages of each triplicate set (3 average values per point), (ii) when distributions are presented, the data from the three replicates were combined into a single dataset (6000 values per curve).Unless otherwise noted, analyses were performed using the VMD/TCL interface or CHARMM.

*Geometric analysis of the lobes*. Three parameters were calculated: (i) the distance between the two COMs of the N-lobe (residues 23 to 105) and the C-lobe (residues 112 to 320), (ii) the angle that the COMs form with the hinge (residue 108), and (iii) the height of ERK was defined as the difference between the maximum and minimum z-component of ERK after alignment using ProDy software [70]. The interaction energy was calculated for two systems: (i) the N- and C-lobe and (ii) T185/Y187 with the kinase. Cluster ensembles were chosen using Chimera and loop figures were based on the representative trajectory [71].

*Loop elongation calculation*. Loop size was evaluated as the maximum distance between the midpoint of the activation loop edges (Cαs of residues 167 and 190) and any other Cα on the loop perimeter.

Contact maps measured the distance between the Cα atoms of each residue pair. Interaction distances considered salt bridges between positively and negatively charged atoms and hydrogen bonds between hydrogen acceptors and donors. Both contact maps and interaction distances analyses were tested for the following criteria: residue pair with an average distance greater than 10 Å (3.5 Å for hydrogen bond) or with a standard deviation greater than 50% of the average were eliminated to ensure that only residues with consistent interactions across all three replicates were included in the results.

*Calculation of phosphorylation site orientation*. Orientation was evaluated by measuring the angle formed between the sidechain oxygen (O_side_) of T185/Y187, its corresponding Cα atom, and the COM of the αF-helix (residues 208-215). The threshold distinguishing between a hindered and an exposed orientation was set to 90°.

## Acknowledgements

This project has been funded in whole or in part with federal funds from the National Cancer Institute, National Institutes of Health, under contract HHSN261201500003I. The content of this publication does not necessarily reflect the views or policies of the Department of Health and Human Services, nor does mention of trade names, commercial products, or organizations imply endorsement by the U.S. Government. This Research was supported [in part] by the Intramural Research Program of the NIH, National Cancer Institute, Center for Cancer Research. The calculations had been performed using the high-performance computational facilities of the Biowulf PC/Linux cluster at the National Institutes of Health, Bethesda, MD (https://hpc.nih.gov/).

