## Supporting Information for "ERK Allosteric Activation: The Importance of Two Ordered Phosphorylation Events"

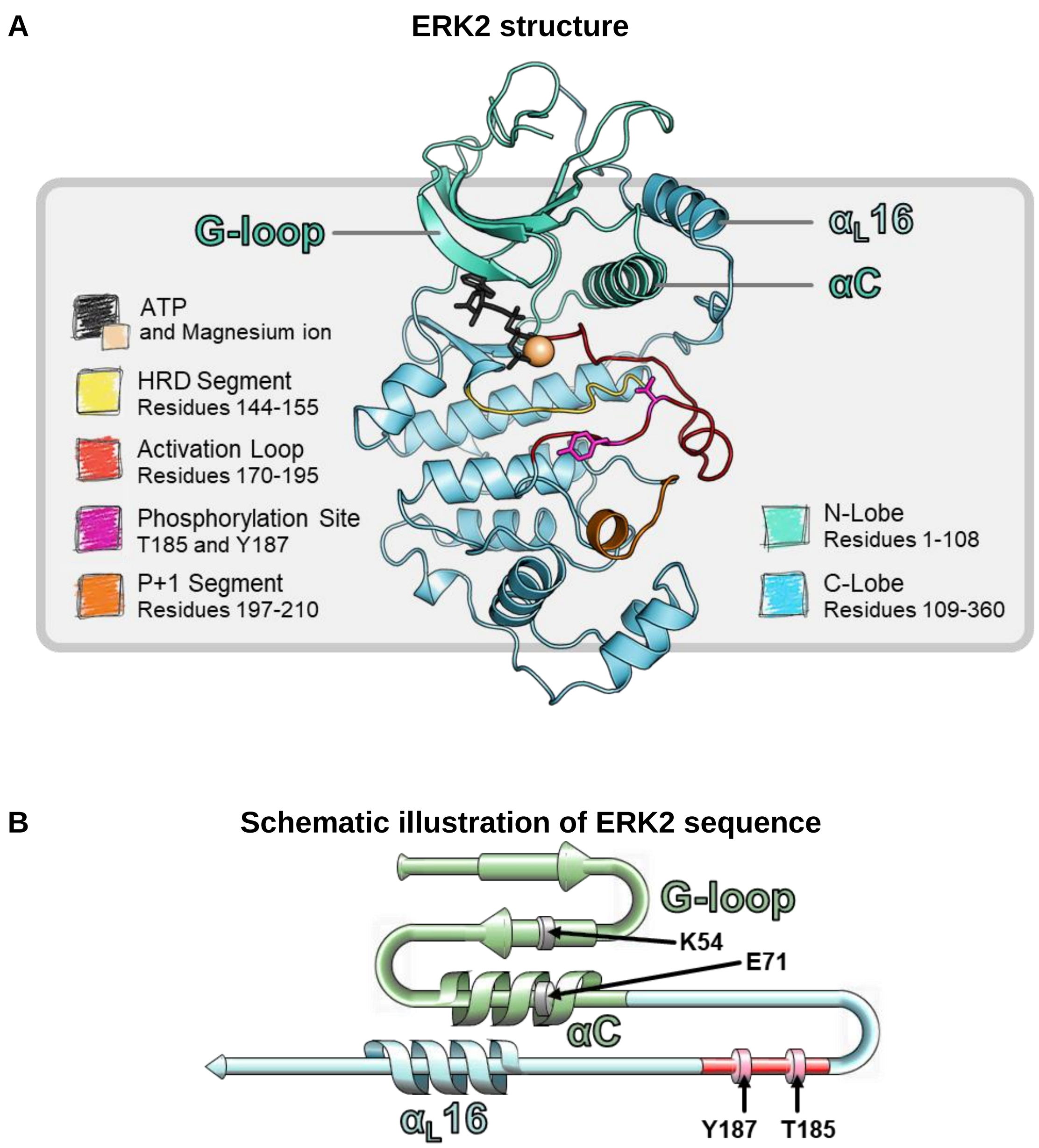


**Figure S1.** Key structural features of ERK kinase. The kinase domain consists of an N-lobe (teal/green) and a C-lobe (cyan). (A) The ATP (black) binding site is located at the interface of the lobes. Above the ATP pocket, the G-loop, formed by a flexible β-sheet, shielding the ATP. Adjacent to the ATP binding site is the regulatory αC-helix, interacting with the C-terminal helix ($\alpha_L$16-helix). The activation loop (red) contains the activation motif TxY (pink). This image is based on the active conformation of ERK2, model $ERK^{p(TY)}$. (B) A sequence illustration highlights important regions of ERK2, including the salt bridge between the G-loop's β-strand and αC-helix, as well as the spatial proximity of αC-helix and $\alpha_L$16-helix.

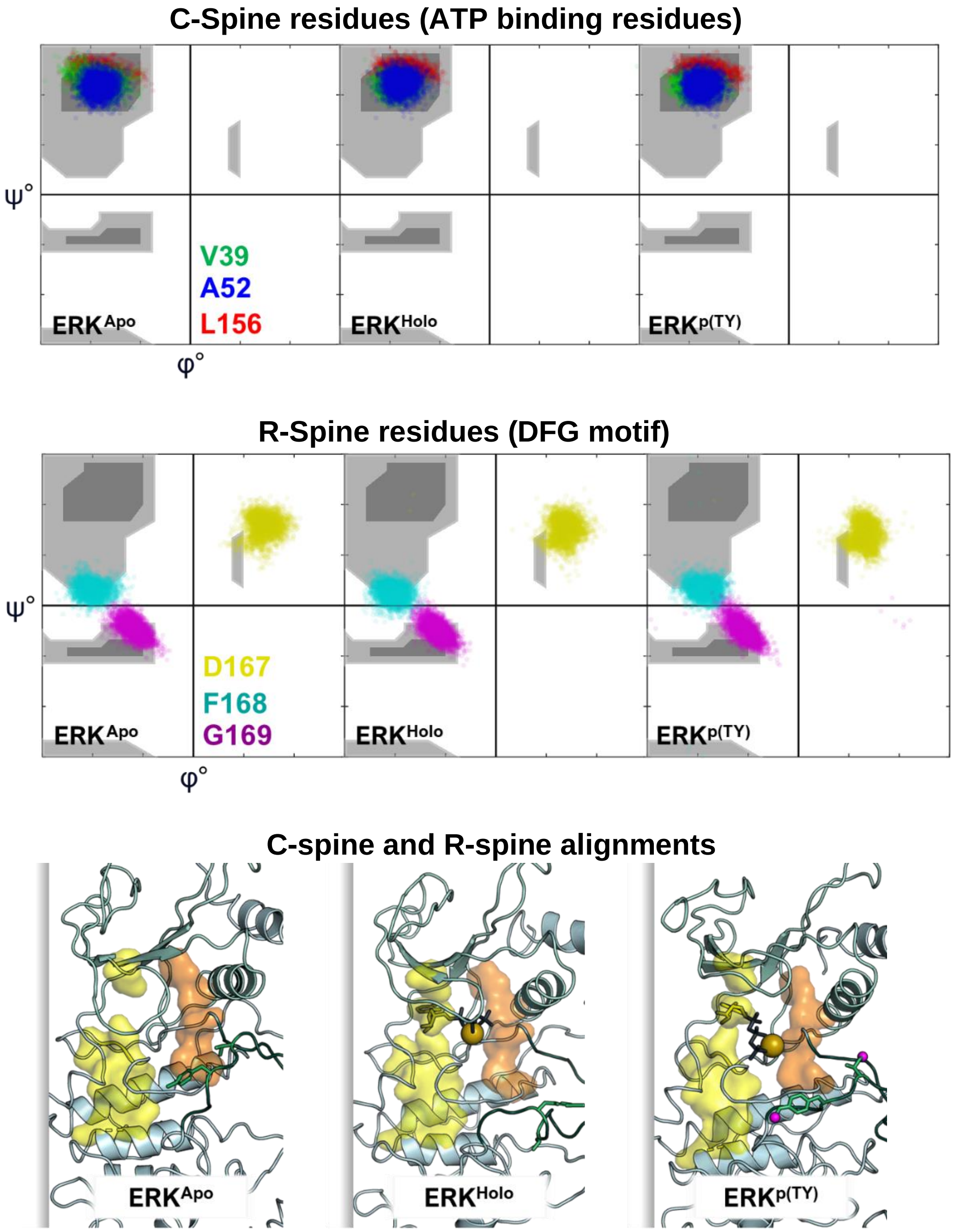


**Figure S2.** The R-spine and C-spine alignments suggest that compactness relies solely on the C-spine. Ramachandran plots of ϕ and ψ backbone dihedral angles for key hydrophobic residues V39, A52, and L156 within the ATP binding pocket that stabilize the C-spine (*top panel*). Ramachandran plots of ϕ and ψ backbone dihedral angles for the DFG motif residues D167, F168, and G169, where F168 stabilizes the R-spine (*middle panel*). Representative snapshots highlighting the R-spine (orange) and C-spine (yellow) for the inactive $ERK^{Apo}$ and $ERK^{Holo}$ and the active $ERK^{p(TY)}$ (*bottom panel*).

### Activation loop elongation and area

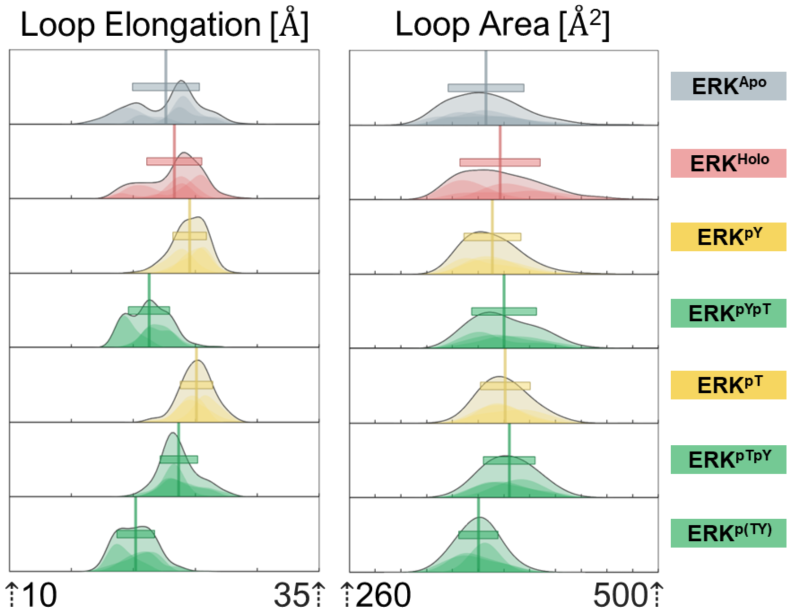


### Schematic illustration of loop area calculation process

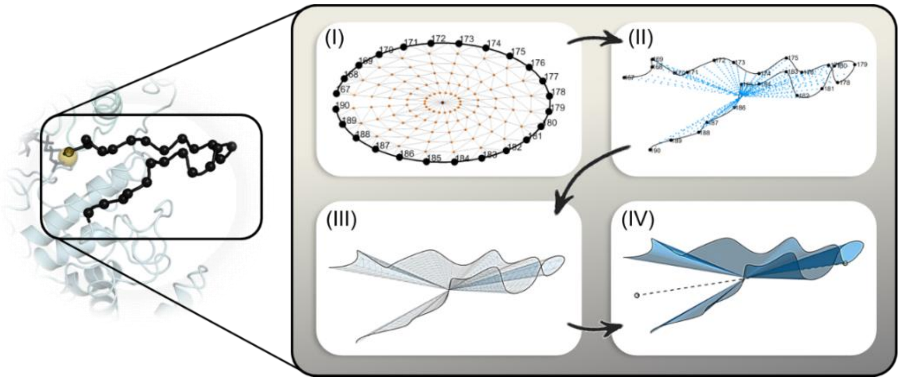


**Figure S3.** Distributions of the activation loop elongation and area (*top panels*). Mono phosphorylation induces allosteric elongations of the activation loop opposite to the compact conformation of the activation loop induced by dual phosphorylation, while all systems show similar loop area values. Schematic illustration of loop area calculation as follows (*bottom panel*): (i) A simplified polygon map of a two-dimensional loop's analogous containing 5 concentric intermediate point series (in orange). (ii) Mapping the perimeter of the activation loop (black) and adding additional geometric center and intermediate concentric points (blue). (iii) Applying the triangular polygon map from the first step to the three-dimensional model (polygons are marked in gray). (iv) Summing the polygons' area to obtain the entire loop surface area.

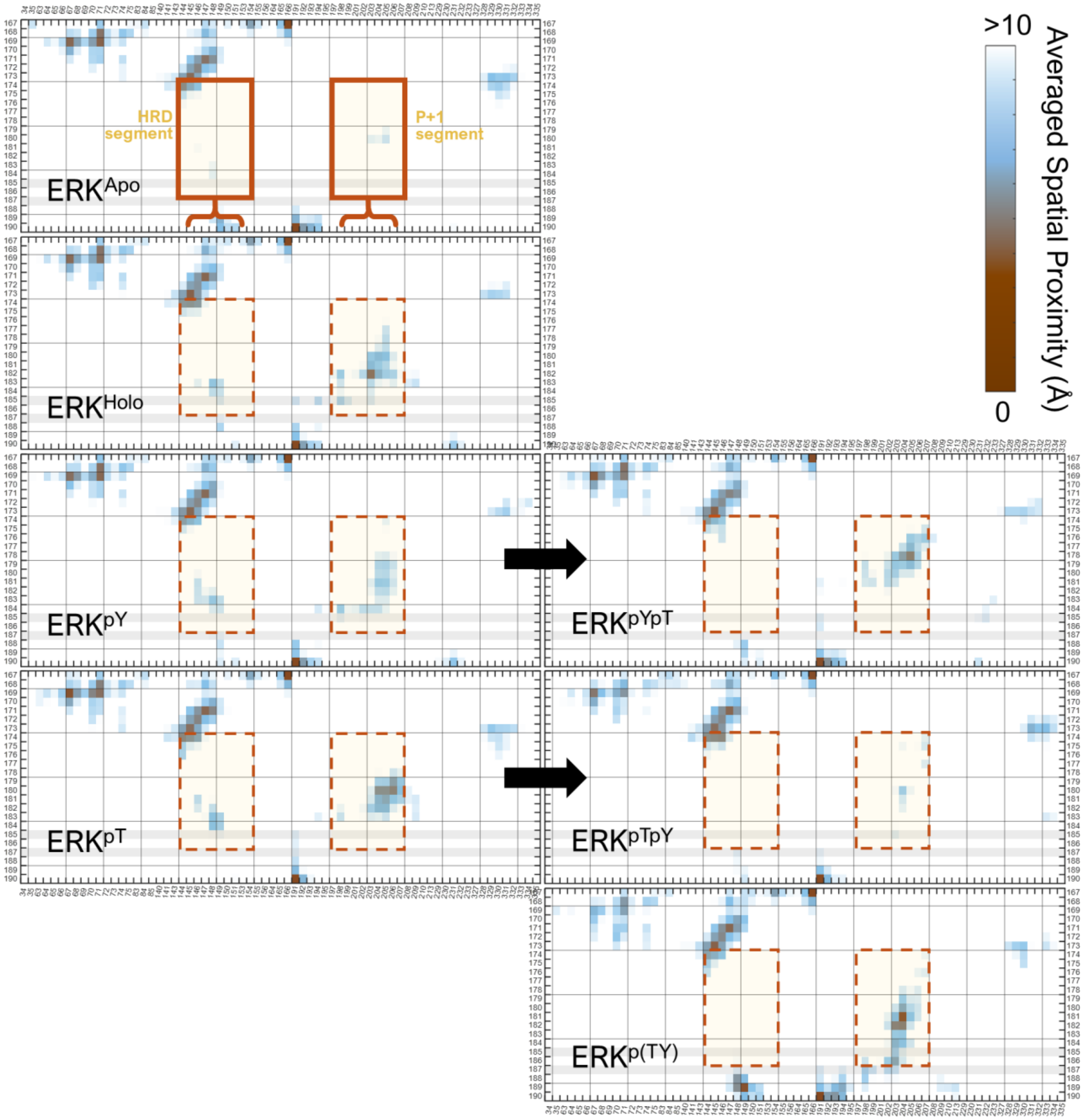


**Figure S4.** Residue-residue contacts for ERK. Contact maps for the activation loop (residues 167-190, y-axis) versus the rest of the kinase domain (x-axis). Spatial distances were computed between Cαs within each residue pair. A threshold of 10 Å and 50% distance and standard deviation were applied, respectively. Residue pairs not meeting these criteria were removed from the maps. The contact maps highlight two regions (along the x-axis) exhibiting significant changes in contacts: the region containing HRD and the P+1 loop segment. See Figure S1 for the ERK structure.

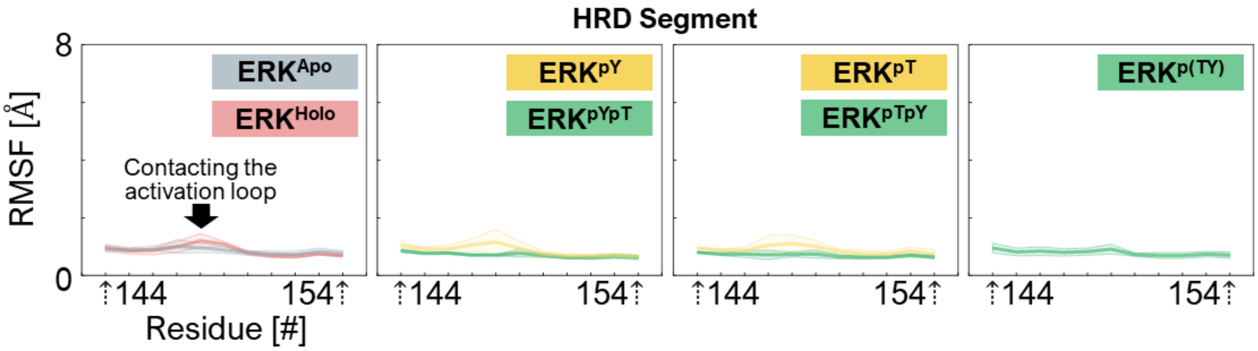


**Figure S5.** Impaired stability of HRD during activation of ERK. Root mean square fluctuation (RMSF) of the region containing HRD (HRD segment, residues 144-154) for the inactive systems in the absence and presence of ATP ($ERK^{Apo}$ and $ERK^{Holo}$), the monophosphorylated systems ($ERK^{pT}$ and $ERK^{pY}$), and the dual phosphorylated systems ($ERK^{pYpT}$, $ERK^{pTpY}$, and $ERK^{p(TY)}$). The average value per residue is plotted as a thick solid line and the standard deviation is illustrated as its faded background color.

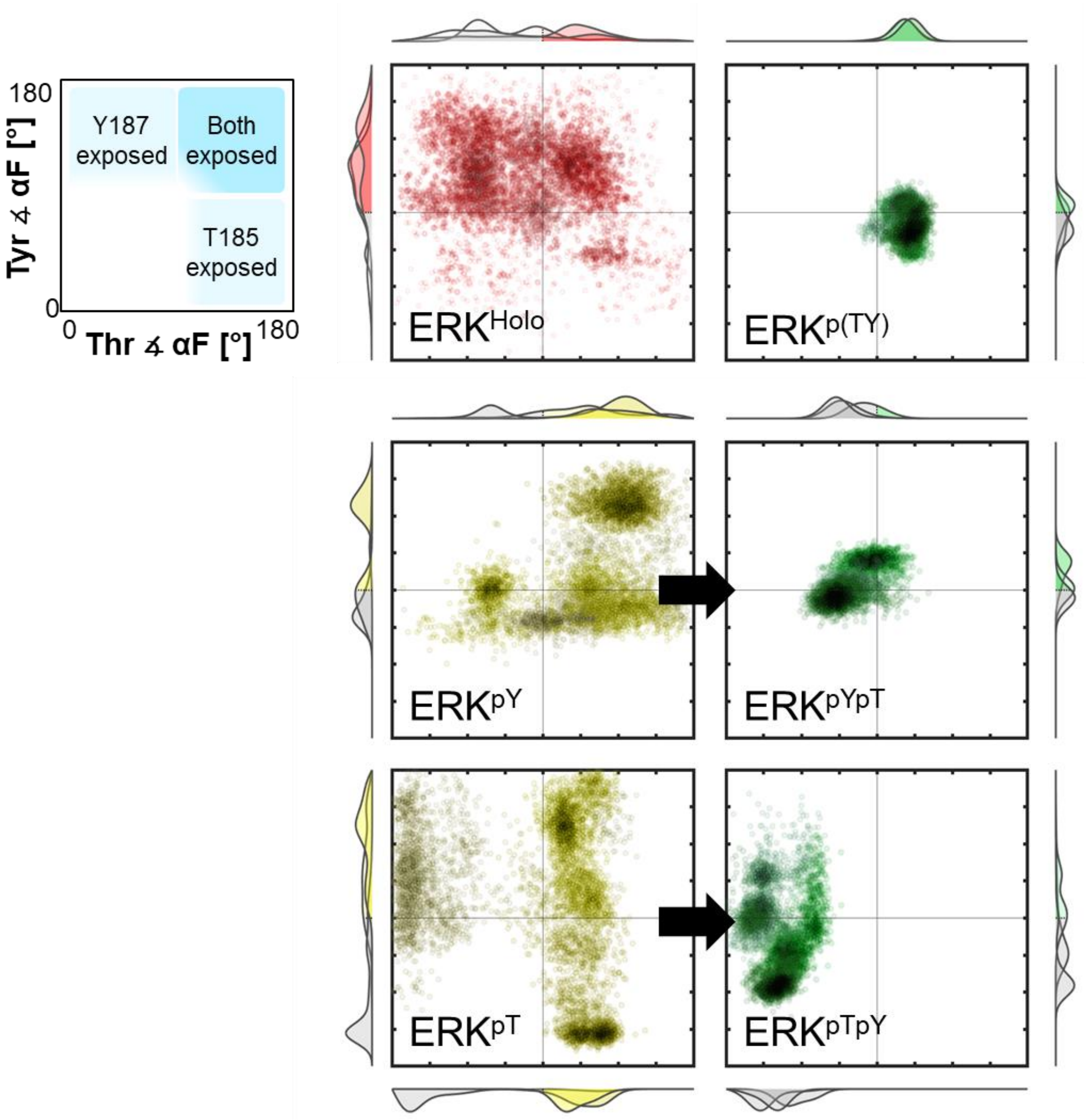


**Figure S6.** Sidechain orientations of T185 and Y187 depending on phosphorylation codes. Two-dimensional (2D) scatter plots representing the orientation angles of the sidechains of T185 (x-axis) and Y187 (y-axis) with respect to the αF-helix for the inactive $ERK^{Holo}$ (red), the monophosphorylated $ERK^{pY}$ and $ERK^{pT}$ (yellow), and the active $ERK^{p(TY)}$, $ERK^{pYpT}$, and $ERK^{pTpY}$ (green). A schematic representation of the orientation plot is shown (*left panel*). The data points indicate the angles formed between the hydroxyl group, its corresponding Cα atom, and the center of mass of the αF-helix (residues 208-215). Angles below 90° indicate potential steric hindrance with the hydroxyl group facing the αF-helix. Angles above 90° indicate enhanced accessibility with the hydroxyl group facing the solvent.

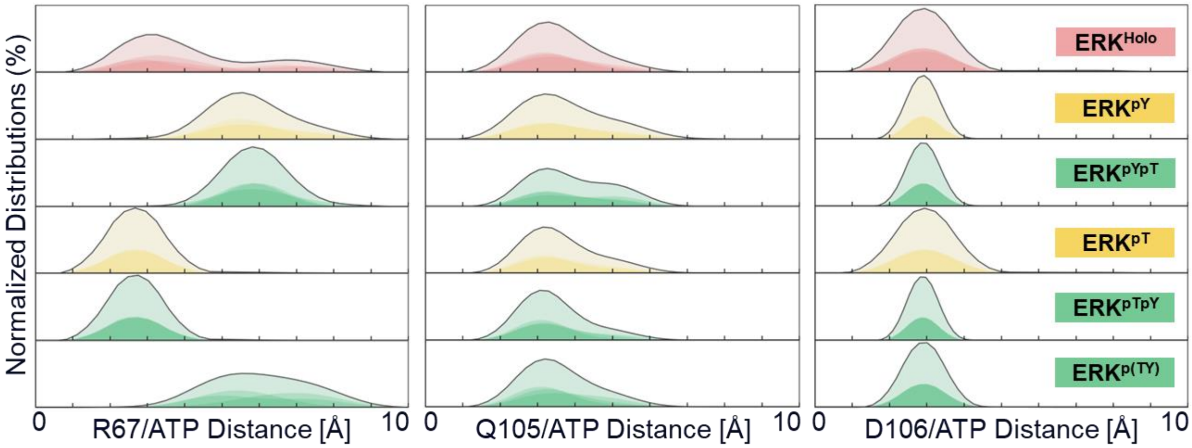


**Figure S7.** Interaction of ATP in its pocket. Distributions of normalized distances from ATP to R67, Q105, and D106. When Y185 is first phosphorylated, the distributions of R67 for $ERK^{pY}$ and $ERK^{pYpT}$ align with the active conformation $ERK^{p(TY)}$. When T185 is phosphorylated first, the distributions of R67 for $ERK^{pT}$ and $ERK^{pTpY}$ correlate solely with the inactive conformation $ERK^{Holo}$. In contrast, Q105 and D106 exhibit similar distance distribution patterns in both inactive and active states.

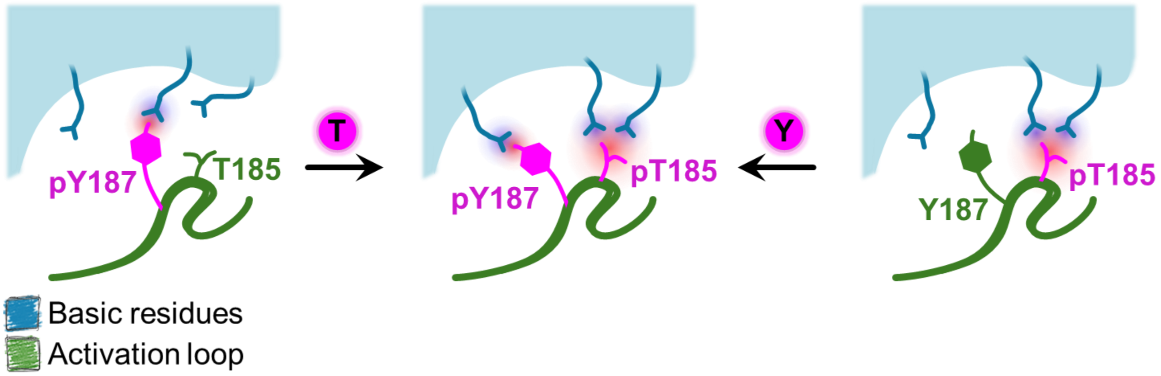


**Figure S8.** Schematic illustration of the dynamic salt bridge formation upon phosphorylation. The weaker interaction pairing of tyrosine may change upon threonine phosphorylation.

**Table S1. Summary of simulated ERK systems.**

| Phosphorylation code | | Non | | Mono | | Dual | | |
|---|---|---|---|---|---|---|---|---|
| State of system | | Inactive | | | | Intermediate | | Active |
| **System** | | **ERK$^{Apo}$** | **ERK$^{Holo}$** | **ERK$^{pY}$** | **ERK$^{pT}$** | **ERK$^{pYpT}$** | **ERK$^{pTpY}$** | **ERK$^{p(TY)}$** |
| **Components** | ATP, $Mg^{2+}$ | - | ✓ | ✓ | ✓ | ✓ | ✓ | ✓ |
| | pT185 | - | - | - | ✓ | ✓ | ✓ | ✓ |
| | pY187 | - | - | ✓ | - | ✓ | ✓ | ✓ |
| **Simulation time (µs)** | | 3 µs | | | | | | |
| **Replicates (runs)** | | 3 per system | | | | | | |
| **Trajectories (#)** | | 1 per ns, total of 6000 per replicate | | | | | | |
| **Based on PDB** | | 4IZ5 | | | | *pY | *pT | 6OPG |

* After 3 µs of simulation and convergence of RMSD.

**Table S2. Clustering of activation loop conformations and occurrence.**

| (%) | System | | | | | | |
|---|---|---|---|---|---|---|---|
| Cluster (#) | $ERK^{Apo}$ | $ERK^{Holo}$ | $ERK^{pY}$ | $ERK^{pYpT}$ | $ERK^{pT}$ | $ERK^{pTpY}$ | $ERK^{p(TY)}$ |
| 1 | 10.8 | 24.0 | 22.5 | 24.0 | 19.8 | 25.1 | 27.3 |
| 2 | 8.7 | 9.0 | 16.0 | 17.0 | 11.6 | 7.2 | 23.8 |
| 3 | 8.2 | 6.2 | 7.8 | 8.0 | 9.3 | 5.7 | 12.0 |
| 4 | 7.0 | 5.5 | 6.5 | 6.5 | 9.0 | 5.0 | 8.3 |
| 5 | 6.5 | 5.5 | 5.8 | 6.0 | 6.7 | 4.7 | 3.5 |
| 6 | 5.0 | 4.2 | 5.5 | 5.0 | 5.2 | 3.8 | 2.5 |
| 7 | 4.5 | 3.7 | 4.0 | 2.7 | 3.7 | 3.7 | 2.3 |
| 8 | 3.8 | 3.2 | 3.5 | 2.5 | 2.7 | 3.3 | 2.2 |

Occurrence of each cluster during the last 2 μs of the simulation trajectories.

**Table S3. Occurrence of residues interacting with ATP.**

| (%) | System | | | | | | ATP binding atom | *Interaction type |
|---|---|---|---|---|---|---|---|---|
| Residue | $ERK^{Holo}$ | $ERK^{pY}$ | $ERK^{pYpT}$ | $ERK^{pT}$ | $ERK^{pTpY}$ | $ERK^{p(TY)}$ | | |
| Q105 (side) | 63 | 57±3 | 48 | 60 | 68 | 62.1 | $NH_2$ | H-Bond |
| D106 (bb) | 100 | 100 | 100 | 100 | 100 | 100 | $NH_2$ | H-Bond |
| S153 (bb,side) | 4.7 | 0.2 | 3.3 | 5.4 | 1.0 | 25.1 | $O^{2'}$ $O^{3'}$ | H-Bond |
| D111 (side) | 19.9 | 0.9 | 3.6 | 31.6 | 6.6 | 2.9 | $O^{2'}$ $O^{3'}$ | H-Bond |
| A35 (bb) | 77.4 | 37.1 | 5.6 | 7.6 | 46.7 | 38 | $O^{\beta}$ | H-Bond |
| Y36 (bb,side) | 0.1 | 0.8 | 0 | 2.6 | 9.4 | 0 | $O^{\beta}$ | H-Bond |
| K54 (side) | 100 | 100 | 100 | 100 | 100 | 100 | $O^{\beta}$ $O^{\alpha}$ | Electrostatic |
| R67 (side) | 94±9 | 96±8 | 99±2 | 100 | 100 | 92±6 | $O^{\gamma}$ | Electrostatic |
| K151 (side) | 92±5 | 96±7 | 100 | 100 | 78±18 | 100 | $O^{\gamma}$ | Electrostatic |

Occurrence of interaction noted as average ± STD (red, orange and yellow markers: STD greater than 50%, 20%, 10% from the average per model). bb=backbone, side=sidechain.

*Occurrence threshold of interaction: < 10 Å for salt bridges, < 3.5 Å for hydrogen bonds.

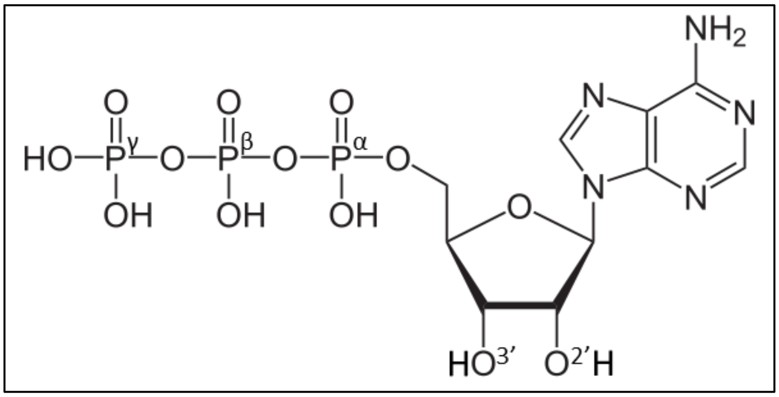


**ATP binding atom legend**
